# Collective motion of bacteria promotes soil water transport

**DOI:** 10.64898/2026.05.20.725210

**Authors:** Beatriz Meza-Manzaneque, Emma Gomez-Peral, Gloria de las Heras-Martinez, Iker Martin-Sanchez, Nicola Stanley Wall, Benjamin Perez-Estay, Anke Lindner, Eric Clement, Natalia Elguezabal, Lionel X. Dupuy

## Abstract

Although rhizosphere microbiomes are known to enhance plants’ resistance to water stress, it is believed that only fungi actively contribute to the transport and uptake of water. We investigated the biomechanical impact of bacterial motility on water transport in soil by combining surface tension measurements and water infiltration experiments in soil microcosms. We observed that flagellar-based motility in the of the rhizobacteria *Bacillus subtilis* cells reduces the apparent surface tension of fluids by up to 15%. The effect reported depends on cell density and swimming speed, confirming its biomechanical origin, and was able to accelerate water infiltration and rewetting of soil. We conclude that Bacillus subtilis facilitates soil water transport through the deformation of air water interfaces in pores.

**Significance:** Water and light limitations to photosynthesis rarely occur simultaneously enabling plants in arid environments to allocate a greater proportion of assimilated carbon to belowground growth, particularly to rhizodeposition. Using microbial activity to convert chemical energy into mechanical work within soil pores offers a major opportunity for improving water use efficiency in agriculture, especially as farming shifts from polluting, energy-intensive mineral fertilisers toward resilient biological fertilisation alternatives.

## Introduction

A motile Escherichia coli bacterium exerts a force dipole of typically 10⁻¹ J on water, which corresponds to the amount of work needed for the bacterium to swim over its own size (1,2). This energy is produced internally via flagellar rotation and for a bacterial swimming speed of 20 microns per second and a bacteria suspension at a cell density of 10^8^ ml⁻¹, the power required to sustain this activity is approximately 10⁻ W ml⁻¹ meaning that over the course of a day, the energy budget is equivalent to the raise of 1 ml of water by approximately 1 cm. Yet, all recent studies showing the effect of rhizosphere microbiota on drought resistance in plants attribute these effects to the presence of fungi which abundance is increased during the water deficit (3). It was proposed that, like with phosphorus uptake, fungal species may form associations with plant roots and produce hyphae that allows the root to connect to distant water (4–6).

In the soil physics literature, it is well established that bacterial activity affects many soil physical properties, but for a large part, effects on soil hydraulics have been linked to secretions that modify the physics and chemistry of solutions (7). Extracellular Polymeric Substances (EPS) secreted by soil bacteria are high-molecular-weight hydrophilic compounds that act to slow down soil desiccation. They increase the viscosity of the soil solution, making it dependent on the flow rate (8). They also block or reduce the conductivity of some pores (9). As a result, water becomes more strongly absorbed, and evaporation is delayed. Furthermore, other secretions may facilitate soil rewetting by producing surfactants: *Pseudomonas* produces rhamnolipids (10), *Bacillus* species produce lipopeptides (11), *Rhodococcus* or *Mycobacterium* secrete trehalolipids (12), and *Rhizobium* species produce glycolipids (13).

We expect the influence of cell motility on soil water dynamics to be mechanical in nature, to act within the bulk fluid, and to occur at a different spatial scale. The mobility of peritrichous cells such as *Bacillus subtilis* or *Escherichia coli* generates hydrodynamic flow moving outward from the head-tail direction and inwards towards its side, characteristic of so-called pusher-like swimmers. When bacteria align due to external flows, boundaries or interactions with neighbours, they can exert macroscopic stresses on the surrounding fluid. This was shown to lead to the emergence of collective motion and the spontaneous formation of vortices (14–17). When confined in the small channels of microfluidic devices or within liquid crystals, mobility was observed sustaining currents (18–20). Motility was also linked to increased diffusivity of solutes (20–22), which in a soil, may affect nutrient transport to the plant. Motility also affects the viscosity of fluids in ways that directly depend on the swimming behaviour of the microorganisms. ’Pusher’-like swimmers, such as *Bacillus subtilis* bacteria, create extensile flow along their swim axis, reducing the viscosity of their surroundings (23). “Puller” like swimmers such as *Chlamydomonas reinhardtii*, create contractile flows along their swimming axis that increase the viscosity of their suspension (24).

The effect of microorganism motility on macroscopic water movements in soil is thus highly plausible, though it has yet to be experimentally demonstrated. To establish the existence of this effect, we conducted pendant-drop assays to determine whether the motile activity of the common soil rhizobacterium *Bacillus subtilis* can produce a measurable deformation of air-water interfaces. Biomechanical effects were isolated from chemical effects by separating cells from their secretions prior to analysis and using mutant strains impaired in surfactant secretion, flagellum assembly, or motor function. We then confirmed using model soil experiments that the motility-borne biomechanical effects reported can substantially influence soil water transport.

## Results

### Apparent air-water surface tension is reduced by bacterial motility

To study the effect of the motility of *Bacillus subtilis* on the apparent surface tensions y, systematic pendant-drop experiments (Figure 1A) were performed on secretionl1lfree cell suspensions and bacterial secretions extracted by centrifugation (Figure S1). Varying cell types and cell densities were characterised. The experimental procedure was completed within 30 minutes to reduce the influence of newly produced bacterial secretions. Besides wild-type strains (WT), we used mutants with impaired motility, either lacking flagellar filament (Δ*hag*) or rotary motor (Δ*motB*). Additionally, we used a mutant deficient in the production of surfactin (Δ*srfAA*), the natural surfactant secreted by the bacterium.

**Figure 1.**
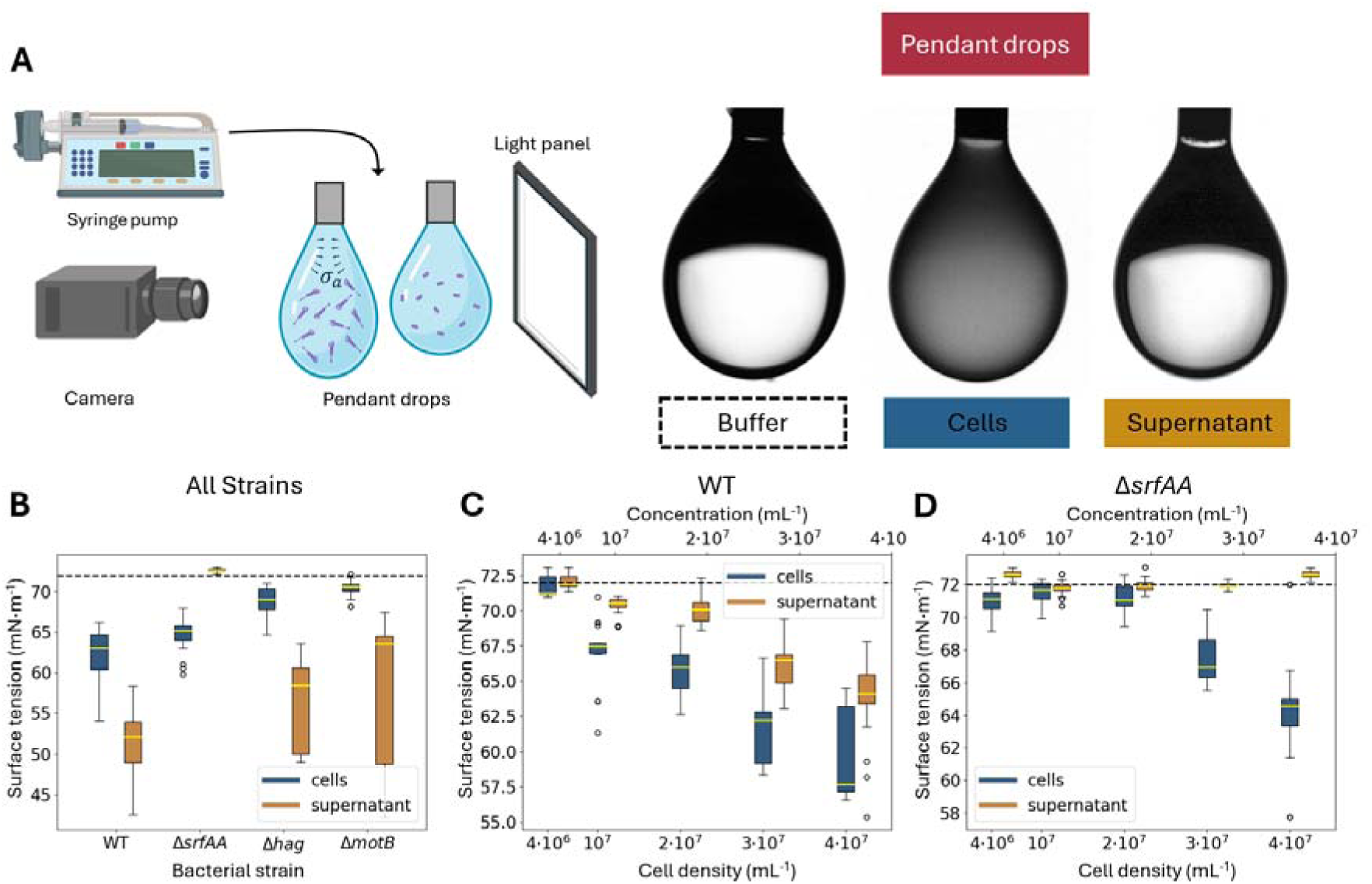
Effect of motility on the apparent surface tension in pendant-drop experiments. (A) experimental setup (left) and shape and appearance of pendant drops for the motility buffer, secretion-free cell suspensions in motility buffer, and supernatant, respectively (right). (B) Apparent surface tension of cell suspensions at a fixed bacterial concentration of 5×10^7^ mL^-1^ (blue), the supernatant resulting from the centrifugation of the cell suspension (orange), and motility buffer (dashed line). (C) Apparent surface tension of the WT bacterial suspension as a function of cell density (blue and bottom axis) and the corresponding supernatant concentration (orange and top axis). (D) Apparent surface tension of Δ*srfAA* bacterial suspension as a function of cell density (blue and bottom axis) and the corresponding supernatant concentration (orange and top axis). Supernatant concentration expressed as a cell density equivalent, see materials and methods “Bacterial strains and growth conditions”.

At a fixed cell density of 5×10^7^ mL^-1^, the supernatants of all bacterial strains expected to synthesise surfactin, i.e., WT, Δ*hag,* and Δ*motB*, showed a consistent reduction of the surface tension to values of between 50 and 60 mN m^-1^ (Figure 1B, orange). WT and Δ*had* supernatants exhibited the lowest values (51.5 ± 0.4 mN m^-1^ and 56.2 ± 0.7 mN m^-1^, respectively). By contrast, the supernatant of a mutation which prevented the synthesis of surfactin (Δ*srfAA*) did not show any reduction in the apparent surface tension (72.6 ± 0.1 mN m^-1^). An analysis of variance showed that the secretion of surfactants had a significant effect on the apparent surface tension (p<0.001). We also looked at the apparent surface tension of cells suspended in a motility buffer (Figure 1B, blue). The apparent surface tension of the immotile bacterial suspensions, Δ*hag* (68.9 ± 0.2 mN m^-1^) and Δ*motB* (70.5 ± 0.1 mN m^-1^) were not drastically different from each other and were similar to the surface tension of the motility buffer, which was equal to that of water (72.8 mN m^-1^). The difference between the suspensions of motile cells was statistically significant, but both WT and Δ*srfAA* consistently exhibited reduced surface tension (62.1 ± 0.4 mN m^-1^ and 64.7 ± 0.2 mN m^-1^) and an analysis of variance showed that motility had a significant effect on the apparent surface tension (p<0.001).

Next, we examined whether the phenomenon observed was dependent on the cell density n. With WT, both the apparent surface tensions of secretions and resuspended solutions decreased with cell density (Figure 1C). The data showed the slope was significantly smaller for secretions than for cell suspensions. In the case of cell suspensions of Δ*srfAA* (Figure 1D), the effect of motility on the apparent surface tension was also density-dependent but non-linear, with effects recorded for cell density above 2×10^7^. No effects of supernatant concentration were observed on the apparent surface tension.

### Reduction of the apparent surface tension is a biomechanical effect

If the reduction in apparent surface tension observed in motile cell suspensions is a biomechanical effect, then the reduction of the surface tension should be directly related to the swimming velocity v in the cell suspensions. To test this hypothesis, the pH of cell suspensions was reduced using hydrochloric acid, and swimming velocity was measured using live microscopy (Figure S1A). Since the proton motive force that actuates the flagella is affected by the pH gradient between the inside and outside of the cell, acidification of the medium is an efficient way to reduce the swimming speed (25), without affecting the surface tension of the solution. Reducing the pH from 6.85 to 5.57 was sufficient to reduce and eventually stop cell motility (Figure S1B). The mean velocity changed from 36.6 ± 1.3 µm s^-1^ to 11.0 ± 0.9 µm·s^-1^, 7.2 ± 0.6 µm s^-1^ and 4.2 ± 0.1 µm s^-1^, respectively, for pH adjusted at 6.85, 6.42, 6.60 and 6.21. We did not observe major shifts in cell counts at these different pH values. However, at pH 5.57, cells sedimented on the surface of the glass slide, demonstrating the absence of motility. In response to changes in pH, the apparent surface tension was observed to rise progressively to values similar to that of the motility buffer (Figure 2A), clearly indicating that the decrease in surface tension is linked to bacterial motility.

**Figure 2.**
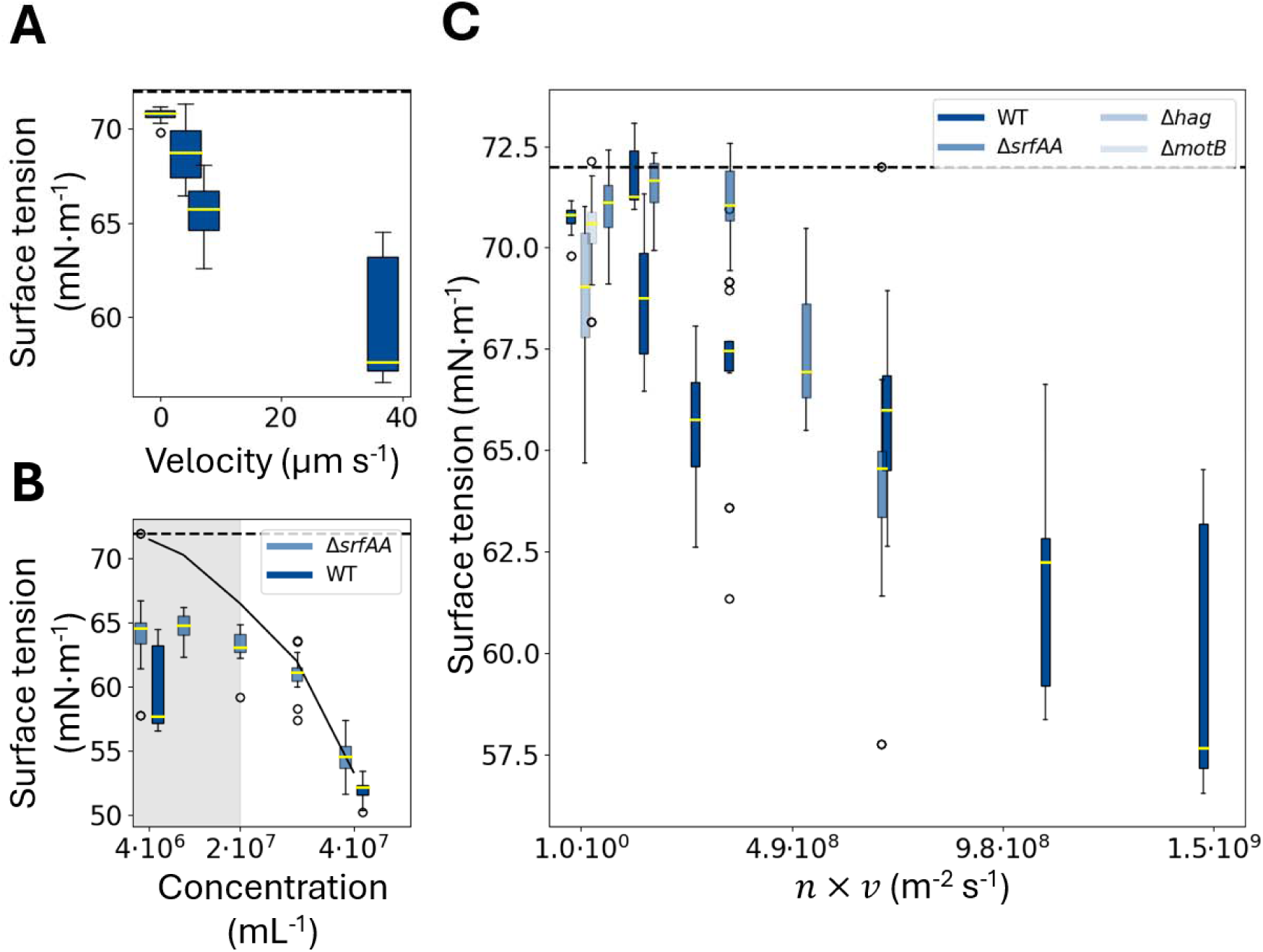
The effect of cell motility on the apparent surface tension is a biomechanical effect. (A) The relationship between bacterial swimming speed and apparent surface tension, obtained using cell suspensions at a density of 4x 10^7^ mL^-1^. The dashed line represents the surface tension of the motility buffer. (B) Effect of the concentration of WT secretions in the solution of Δ*srfAA* cell suspensions maintained at a density of 4x 10^7^ mL^-1^. WT (dark blue) and Δ*srfAA* cells resuspended in WT supernatant at different concentrations (light blue). Apparent surface tension of WT supernatant alone obtained from figure 2B (plain line), and motility buffer (dashed line). Supernatant concentration expressed as a cell density equivalent (mL^-1^), see materials and methods “Bacterial strains and growth conditions”. (C) All experiments conducted on cell suspensions (absence of supernatant) converge toward a single trend relating apparent surface tension to motile activity nx v.

We also examined potential interactions between motility and surfactant production, to ensure that any trace surfactant formed during celll1lsuspension preparation did not interact with cell motility to reduce the apparent surface tension. Δ*srfAA* strains were resuspended in WT supernatant diluted to various concentrations in the motility buffer. At low supernatant concentration, the WT supernatant reduced the apparent surface tension of Δ*srfAA* cell suspensions (Figure 2B, blue), but the reduction was less pronounced than if only supernatant was present (Figure 2B, plain line). However, when more than 50% of the secretions of the WT strains were present, the apparent surface tension of the suspension matched that of the secretion alone. The response of the apparent surface tension to the addition of supernatant was 10 times stronger at high concentration than that obtained at a lower concentration (−3.0 x 10^-6^ mN mL m^-1^ and - 3.2 x 10^-7^ mN mL m^-1^ respectively). When the Δ*srfAA* strains were resuspended in the pure supernatant extracted from WT strains, the apparent surface tension was restored to values similar to those of the WT strains (52.1 ± 0.1 mN m^-1^ and 54.7 ± 0.3 mN m^-1^ for WT and Δ*srfAA,* respectively). We can therefore discard the hypothesis that the effect of motility on the apparent surface tension was due to an interaction between motility and surfactants present at low concentration. The results also show that the chemical and mechanical effects of microbial activity are not cumulative. At low supernatant concentrations, motility in the suspension determined the apparent surface tension measured, whereas at high concentrations the apparent surface tension was governed by surfactant concentration in the solution (Figure 2B).

To further demonstrate that the influence of bacterial motility is of biomechanical origin, we studied the relationship between motile activity and apparent surface tension across all experimental conditions. We chose to represent this activity in the form of the activity flux J_A_ =n x v defined earlier to account for the enhanced diffusivity of a passive tracer (22). In our experiments, the swimming speed v (Figure S1, mm s^-1^) varied as a function of pH and strain type, and the cell density n (Figure 2, mL^-1^) changed according to the dilution level. When representing the apparent surface tension as a function of the active flux for all strains and experimental conditions studied in this paper, the results collapse onto a single curve (Figure 2C) decreasing as expected with the active flux.

### Implications for soil water transport

Soil microbial activity is a critical component of the plant nutrition system and recognising that cell motility could contribute to water transport and ultimately root water uptake is of great significance for agricultural and ecological sciences. We conducted a series of soil water infiltration experiments in a model soil system to assess how motility may also influence water movement when motile cells are held in soil pores. In the absence of bacteria, tracer dye solutions used in the assay were not seen spreading through a hydrophobic dry soil layer even after 144 hours, and there was no water infiltration into the bottom soil layers (Figure 3A, left). In the presence of non-motile bacterial cells (Δ*hag*) resuspended in the tracer dye solution, a significantly increased rate of infiltration and rewetting was observed. After 120 h, the tracer dye was observed consistently at the bottom layer of the microcosms, and by the end of the experiment, the dry soil was entirely stained (Figure 3A, right). This outcome aligns with expectations, since the surfactin produced by bacterial cells reduced the pressure required for pore water entry. When motile bacterial cells were resuspended in the tracer dye, the tracer dye penetrated the dry soil layer even quicker, appearing as early as 24 h after inoculation and 24 h earlier than when non-motile bacteria were used in the microcosm.

**Figure 3.**
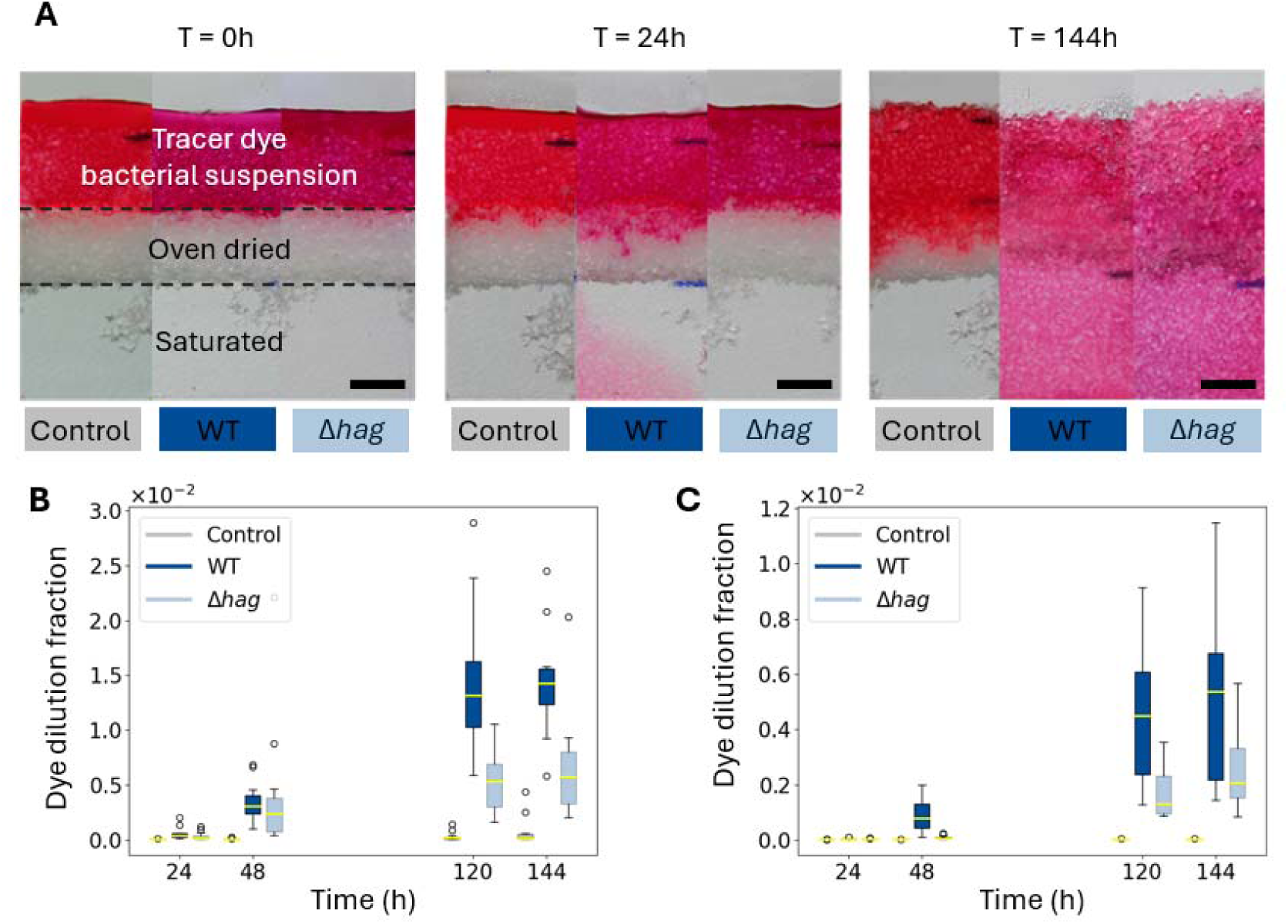
Bacterial motility enhances water infiltration. (A) Dye tracing experiment in an artificial soil microcosm quantifying the infiltration of water through a dry soil layer (scale bar equals 0.5 cm). Three cases were considered: control samples, in which 0.2 mL of the dye solution was added (left); WT samples, in which 0.2 mL of the dye solution containing a WT suspension at n = 2×10^7^ mL⁻¹ was added (middle); and Δ*hag* samples, in which 0.2 mL of the dye solution containing a Δhag suspension at n = 2×10^7^ mL⁻¹ was added (right). (B) Dye dilution fraction f in the dry transparent soil layer. (C) Dye dilution fraction f in the bottom transparent soil layer.

Since immotile cells exhibit a lower infiltration rate than the wildl1ltype strains, it can be concluded that the mechanical energy generated by cell motility enhanced the movement of the soil solution through the dry soil layer below. Even though the experiments lasted long enough for surfactants to start accumulating in the microcosms, we found no evidence that these may have interacted with the motile activity to enhance the infiltration of WT strains (Figure 3B). However, interaction between cell motility and soil structure may have influenced water transport and infiltration. Live fluorescence imaging of the samples revealed that bacteria rapidly accumulated at the air-water interface, possibly due to aerotaxis and gravity (see Figure S3). This was likely to have dramatically increased the cell density at the wetting front and to have magnified the influence of motile activity on water infiltration.

## Discussion

Motile microorganisms possess the unique ability to convert chemical energy from substrates into mechanical work, which, when transferred to the surrounding fluid, can modify its macroscopic properties. Physicists have extensively studied the unusual dynamics of these so-called active fluids, which display properties such as enhanced solute diffusion, reduced viscosity, and collective microbial motion (17,19,21,26,27). Soils, home to some of the most diverse microbial ecosystems on earth, may be significantly affected by these phenomena. Yet, the topic is virtually unexplored. Current theories of transport processes in soil are still largely grounded in classical frameworks developed from past centuries, including Fick’s laws (diffusion of solutes) (28), the Richards and van Genuchten equations (water transport) (29), the Keller–Segel model of chemotaxis (30), or filtration theory (microbial retention in soil)(31). Critically, none of these models account for the active stresses imparted to the fluid by motile soil microorganisms.

We prove here, for the first time, that in the absence of surfactant compounds, droplets of bacterial suspensions present a decrease in apparent surface tension directly related to bacterial motile activity, and that this observation can be attributed to a shift in the balance of forces within the fluid (biomechanical effect), not resulting from changes in the interfacial tension of water. To understand the forces at the origin of this effect, it is useful to first estimate the order of magnitude of the active stress a_a_ in a bacterial suspension of concentration n. For bacteria aligned parallel to the droplet interface, such active stresses have been shown to exert extensile stresses along the interface (32), potentially influencing measurements of the apparent surface tension. However, the dipole strength of a bacterium such as E coli 10^-18^ J (1), and with the cell density reported here (n= 4 x 10^7^ mL^-1^, the resulting active stress a is of the order of 10^-4^ Pa.

Even with the likely 10 fold increase of the dipole strength of *Bacillus subtilis* suspensions, the active stress generated is not sufficient to explain a reduction of 10 mN m⁻¹ in the apparent surface tension, which corresponds to a force of approximately 10^-4^ N in a drop of liquid with a diameter of 3.3 mm (mean diameter in experiments), and a pressure difference of ≈ 12 Pa. This is four orders of magnitude larger than what the bacterium’s classical active stress can generate.

The effect of motile activity on the apparent surface tension car therefore be attributed to the distribution of cells within the drop of liquid. Arguably, aerotaxis and sedimentation leads to accumulation of cells at interfaces, increasing the local bacterial concentration such that a closely packed bacterial layer may indeed generate the necessary active stresses to deform drops of liquids. Additional effects include an increase in the internal droplet pressure, arising from the recently proposed swim pressure induced by confinement (33) . It has also been shown recently that collective motion can arise at cell densities lower than previously assumed (17). Such macroscopic flows in our experiments may have led to significant viscous stresses at the droplet interface and further changes of its shape. Future livel1limaging studies resolving bacterial accumulation and dynamics within pendent drop assays will be essential for quantifying the forces responsible for the apparent reduction in surface tension reported here.

Beyond this, the discovery has fundamental consequences for agricultural sciences, as polluting and energyl1ldemanding mineral fertilisers are poised to be replaced by biological fertilisation approaches. Critically, our results reveal that the motile activity of rhizobacteria may profoundly influence soil water availability to crops, emphasising the need of integrated approaches to nutrition and fertilisation within sustainable soil management practices. How such integration can be achieved, however, remains uncertain due to the complexity of natural soils. Structures vary widely across soil types, affecting pore size distribution and, in turn, the volume through which collective motility can emerge. Water content determines the degree to which microbes are confined into water films, further channelling the direction of the microbial motion as well as modifying their speed (34). Finally, soil microorganisms sense chemical gradients of oxygen and nutrients originating from plant roots, and these cues can, at times, alter their motility (35). A primary challenge, therefore, is to characterise how these factors influence motile activity in soil and subsequent effects on plant transpiration.

## Materials and methods

### Bacterial strains and growth conditions

Bacterial strains were derived from *Bacillus subtilis* (NCIB 3610) and NRS1473 (NCIB 3610 *sacA*::Phy-spank-*gfpmut2* (kan) (36)) which was modified to constitutively express a green fluorescent protein and was considered as the reference strain (later referred as WT). Other strains studied included NRS6959 (NRS 3610 Δ*hag sacA*::Phy-spank-*gfpmut2* (kan) (37)) which lacked flagellar filament and was referred as Δ*hag*; NRS6963 (NRS 3610 Δ*srf*AA (kan) *amyE*::Phy-spank-*gfpmut2* (cml) (37)) which was unable to produce surfactin (referred as Δ*srfAA*); and NRS3434 (NCIB 3610 Δ*motB* Δ*ywsC*::spc (38) which flagellum is assembled but have loss of motor (referred as Δ*motB*), this last strain has another mutation to restrain γ-PGA production since deleting the *motB* gene is associated with an overproduction of this exopolymer (38). Strains were streaked from stock cultures preserved in glycerol at -80 °C on Luria-Bertani (LB) agar plates containing 10 μg mL^−1^ kanamycin, 100 μg mL^−1^ spectinomycin or 5 μg mL^−1^ chloramphenicol, as required. Bacterial colonies were then inoculated in M9G medium (200 mL M9 salts solution 5x, supplemented with 2 mL of 1M MgSO_4_ solution, 0.1 mL of 1M CaCl_2_ solution, 2 g glucose and 0.5 caramino acids per litre) and incubated while shaking for 20-24 hours at 28°C. The cell density was determined from the optical density (OD) of cell the suspensions using a Den-600 photometer (Biosan, Latvia). Calibration between optical density and cell density were obtained using a flow cytometer (CytoFlex, Beckman Coulter, United States). Bacterial suspensions were centrifuged, and the supernatant was removed and kept for measurements of apparent surface tension. Cells were then resuspended in motility buffer (20 mL of 1M Sodium lactate solution, 2 mL of 1mM L-methionine solution, 2 mL of 100mM EDTA solution and 12.5 g of L-serine per litre) for characterisation of the apparent surface tension.

In a first experiments, all bacterial strains were grown at a cell density of 5x 10^7^ mL^−1^ and the resulting suspensions were characterised for apparent surface tension of supernatant and cell suspensions in motility buffer. In a second step, density dependence bacteria was established with bacterial cultures initially grown at a cell density of 4x 10^7^ mL^−1^ before centrifugation and resuspension in different volumes of motility. In these experiments, dilutions were also applied to the supernatants. Interactions between motility and secretion effects, were characterised by measuring the apparent surface tension of Δ*srf*AA cells resuspended in WT secretions across a range of dilutions. Finally, the effect of cell velocity on the reduction of the apparent surface tension was assessed by adjusting the pH of the cell suspensions in motility buffer using HCl. Throughout the manuscript, both cell density and concentration of the supernatant were expressed as mL^-1^. For the supernatant, the concentration indicated is the equivalent cell density of the suspension from which the supernatant can be obtained (Figure 1 C&D and Figure 2B).

### Measurement of surface tension

Surface tension measurements were performed using the pendant drop method. Drops of suspensions with a volume of approximately 170 µL were obtained using a syringe pump (Pump 11 Pico Plus Elite, Harvard Apparatus, United States), a blunt-end needle with an outer diameter of 0.82 mm (Fisnar QuantX™, Ellsworth Adhesive, Spain), and a back-light screen. Images were acquired with a CMOS camera (DCC1545M, Thorlabs, UK) and a telecentric 1X objective lens (375-036-2, Mitutoyo, Germany). For each single surface tension data point of a given suspension, nine different droplets were imaged. Surface tension values were obtained using a custom program (Python 3) that integrates the Bashforth-Adams formulation of the Young-Laplace equation. The algorithm fits the Bond number, the apical radius, and a vertical apex offset to reproduce the observed drop profile. The apical radius used to normalize the Young-Laplace equation was considered an independent free parameter, although an initial ansatz was obtained by fitting a 4th-order polynomial to the apex. To correct for camera misalignments, the axis of gravity was determined to be the one that maximizes droplet shape symmetry. Finally, the surface tension was obtained from the fitted Bond number, apical radius, gravity, and water density.

### Bacterial velocity

Measurements of swimming speed and direction were obtained on cell suspensions at a bacterial concentration of 4x 10^7^ mL^-1^. A drop of 5 µL of cell suspension was placed between a microscope glass slide (76 x 26 x 1 mm3, VWR, Spain) and a gas-permeable cover slip (25 x 75 mm, Ibidi, Germany), separated by double-sided tape (Arcare 94119, Adhesives Research, Ireland). Live microscopy data were acquired from an inverted microscope (DMi8, Leica, Spain) with a 20x objective at a frame rate of 90 ms for 3.6 seconds (40 images). Trackpy (39) was used to track movement of bacteria in image stacks.

### Microcosm experiment

Water infiltration experiments were performed in model soil systems, that is small volumes of transparent soil contained in a chamber of about 8 cm^3^. Transparent soil was prepared as described earlier by (40). Briefly, Nafion pellets (NR50 1100, Ion Power Inc, USA) were fractured to a texture like sand (0.10 to 1.25 mm) using a freezer mill (6850 Freezer/Mill, SPEX CertiPrep, UK) and a series of sieves. The pH was then adjusted by washing it with Hoagland nutrient solution.

Microcosm chambers were assembled from microscope glass slides (76 mm × 26 mm × 1 mm, VWR, Spain) following the protocol described by (41). Glass slides were bonded with 4 mm polydimethylsiloxane spacers (PDMS, SYLGARD 184, Sigma-Aldrich, Spain) after oxygen plasma surface treatment for 15 s at 100 W (HPT-100, Henniker Plasma, UK), pressure was applied to ensure a strong bond between all surfaces in contact. The bottom of the microcosm was filled with a 1.5 cm layer (1.4 g of dry weight) fully saturated in water (0.2 mL of water). Then, a 0.8 cm layer of oven dried (105 °C, 24 hours) transparent soil was added (approximate dry weight of 0.7 g). The last 1 cm of soil (approximately 0.9 g of dry weight) was also fully saturated with water.

The dye tracing experiment developed by (41) was applied to study how motility affected the infiltration of water through the dry soil layer. Bacterial suspensions were mixed with a food dye (0.1 mL mL^-1^, red food colorant, Vahiné, Spain) and 0.2 mL of the suspension (n=2.0x10^7^ ) added to the top layer of the microcosm chambers, the microcosm chambers were closed with parafilm tape and placed in a growth chamber at 23 °C, 60% humidity. Distilled water was added to the samples using a syringe to maintain the top layer of transparent soil saturated. Images were taken using a Canon EOS 2000D camera (Canon, Spain) to quantify the dilution fraction of the tracer dye f. To visualise the distribution of bacteria in the soil images were also acquired using a custom-made horizontal fluorescence microscope composed of a Retiga R6 CCD camera (Teledyne, UK) and a 1X PLAN objective with 50 mm aperture with 2 mm field of view. Immediately after introducing the dye, and 24, 96, 120 and 144 hours after the first image. The experiment was repeated 3 times with a total of 15 samples with bacteria and 13 controls samples without bacteria. Characterisation of the infiltration of the dry soil layer was performed following the protocol developed by (42). Briefly, images were acquired using a Canon EOS 2000D camera (Canon, Spain) and relationships were established between the dye dilution fraction f and the image saturation value.

### Statistical analysis

Analyses were performed using R (R Core team 3.1.2). The effect of motility and surfactant production was assessed using analysis of variance and density dependence of the phenomenon was assessed using a linear model (lm library).

## Acknowledgments: Funding

We acknowledge the funding from the Spanish Ministry of Science, Innovation and Universities MICIU/AEI/10.13039/501100011033, project PID2020-112950RR-I00 (MICROCROWD).

MICIU/AEI /10.13039/501100011033 and by FEDER, UE, project PID2023-149435OR-I00 (BIOFLOW)

NSW was supported by funding from BBSRC, project BB/Z516600/1.

## Author contributions

Conceptualization: LXD

Methodology: BMM, EGP, GHM, IMS, NSW, BPE, AL, EC, NE, LXD

Investigation: BMM, EGP, BPE Funding acquisition: LXD Project administration: LXD

Supervision: NSW, AL, EC, NE, LXD Writing – original draft: BMM, LXD,

Writing – review & editing: NSW, BPE, AL, EC, NE, LXD

## Competing interests

Authors declare that they have no competing interests.

---

**Figure S1.**
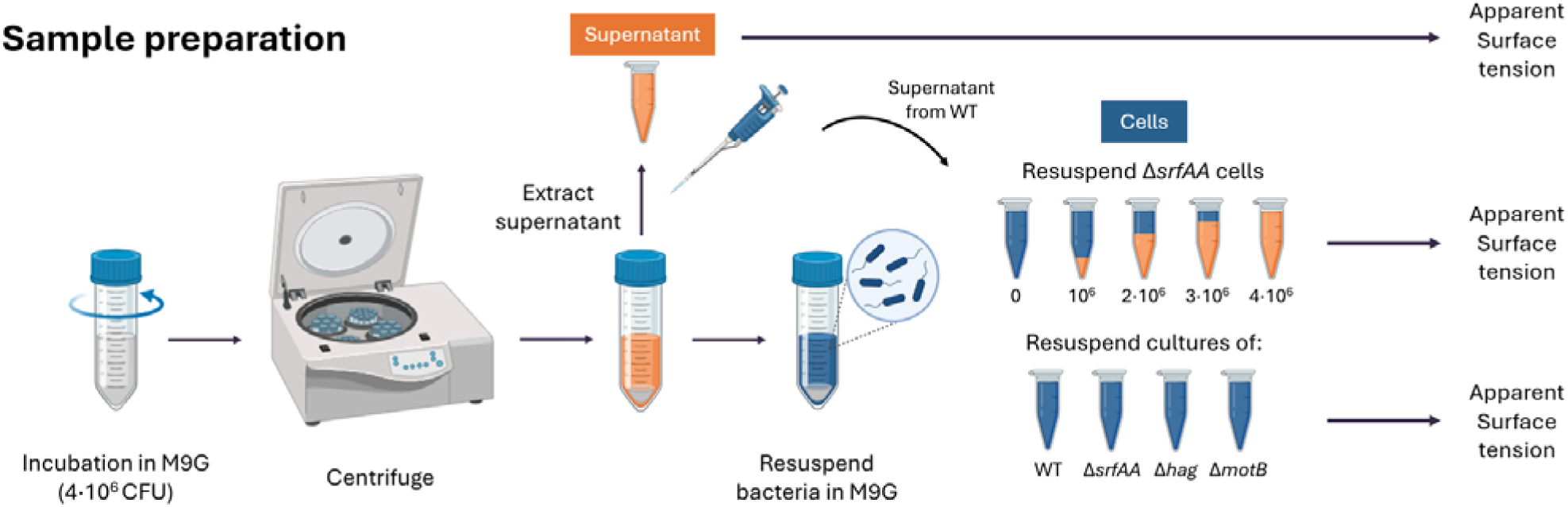
Protocol used for characterising the effect of motile activity on the apparent surface tension of cell suspensions. Cell suspensions are centrifuged, and the supernatants containing secreted compounds were collected. The apparent surface tension was measured for both the cell-free supernatants and for cells that had been resuspended in a motility buffer. For the strain that did not produce surfactin (Δ*srfAA*), cells were also resuspended in the supernatant of WT strains at a range of dilutions.

**Figure S2.**
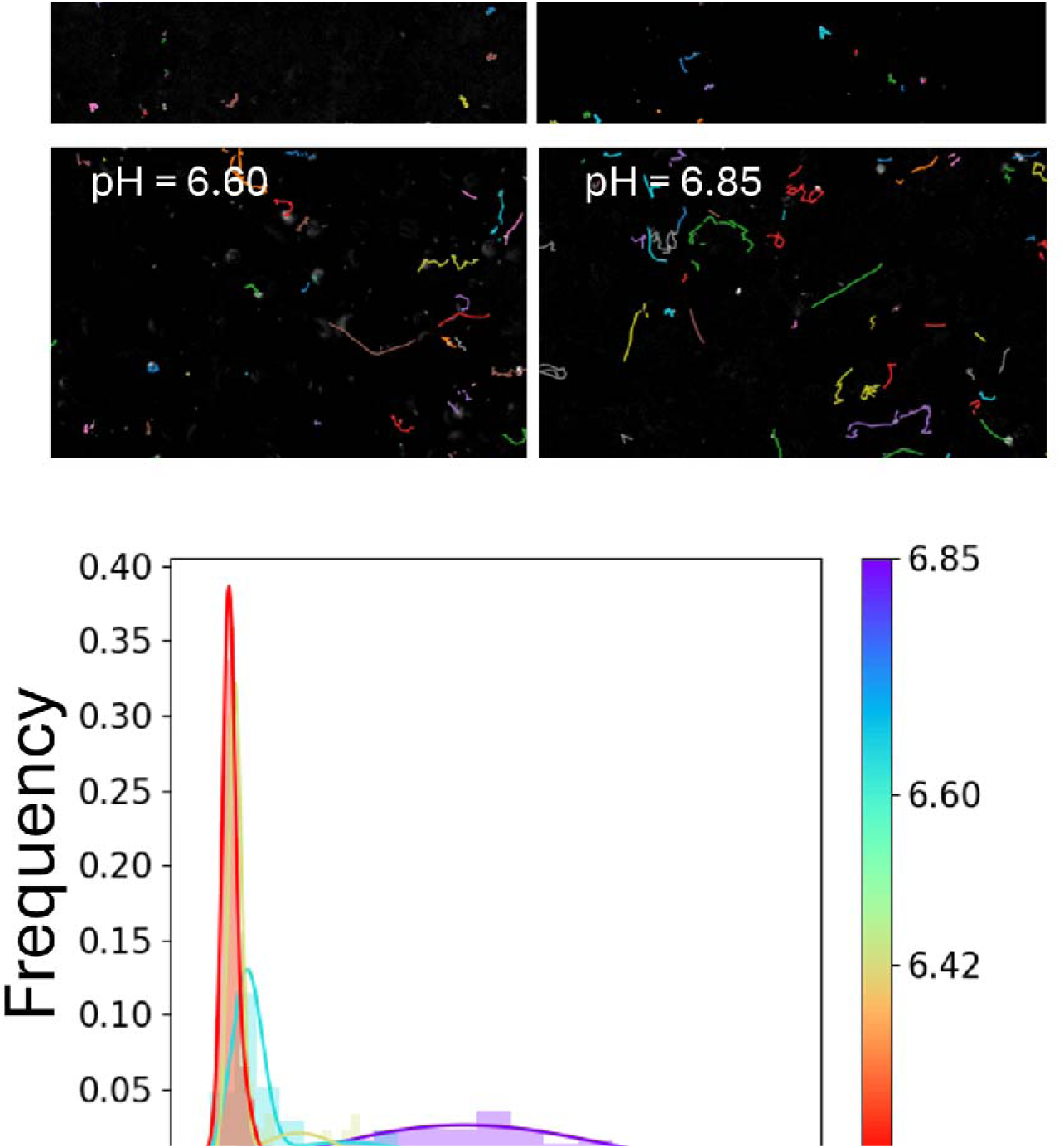
Effect of acidity on cell velocity. (A) Microscope images of WT bacterial suspensions across a pH gradient (from more acidic in the upper left to less acidic in the lower right), with individual cell movement tracked using a particlel1ltracking software. (B) Probability density distribution of swimming speeds at different pH.

**Figure S3.**
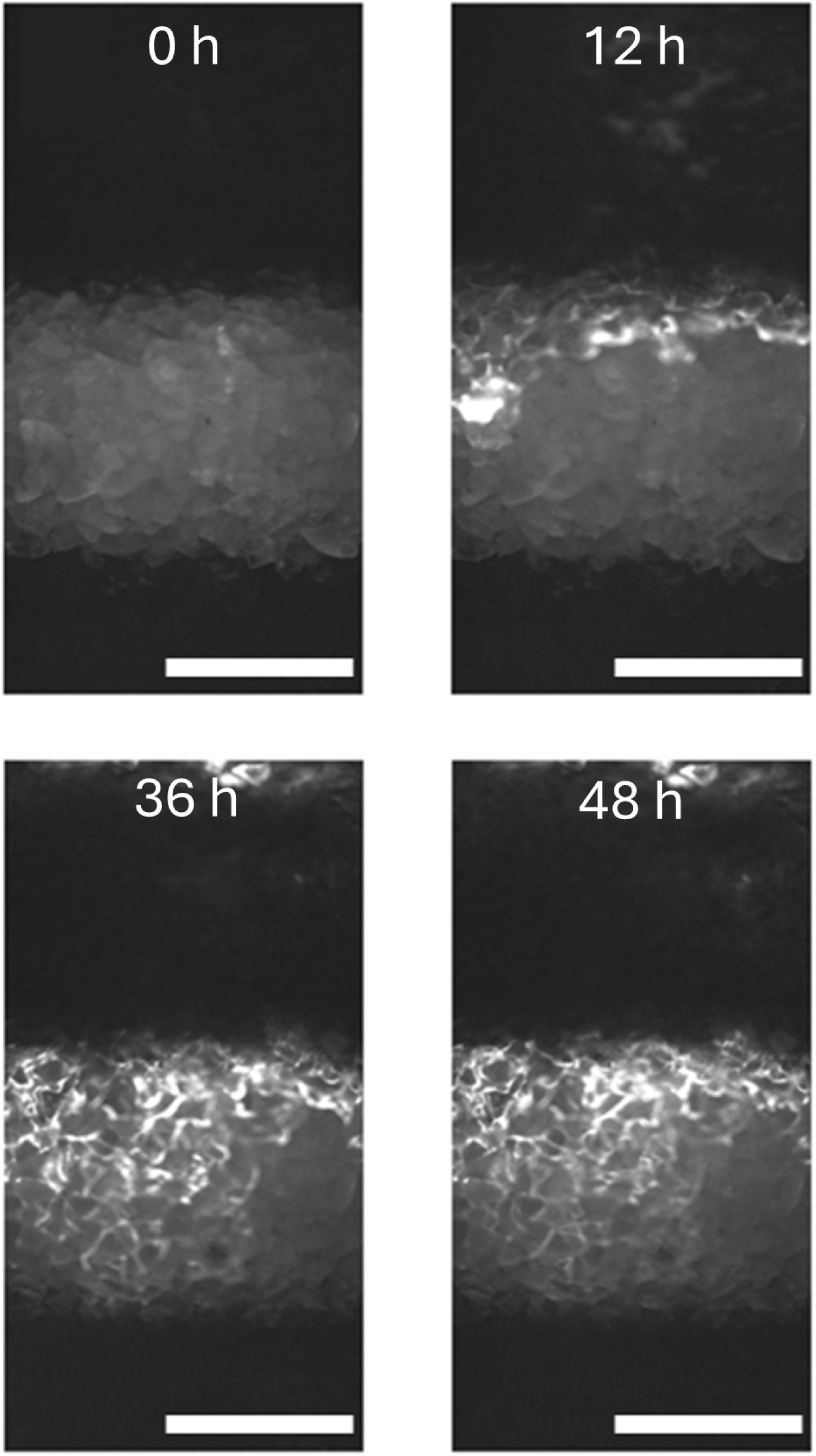
Live fluorescence imaging showing bacterial cell distribution in microcosm.

**Table S1.** List of bacterial strains used in the study.

| Name | Name given in this study | Genotype <sup>a</sup> | Reference / Construction <sup>b</sup> |
| --- | --- | --- | --- |
| NCIB 3610 |  | Wild type prototroph | *Bacillus* genetic stock centre |
| NRS1473 | WT | NCIB 3610 *sacA::Phy-spank-gfp mut2* (kan) | (Verhamme et al., 2007) |
| DS1677 (stocked as NRS2253) | | NCIB 3610 Δ*hag* | (Blair et al., 2008)<br>(kindly provided by Prof Daniel Kearns) |
| NRS6959 | Δ*hag* | NCIB 3610 Δ*hag sacA::Phy-spank-gfp mut2* (kan) | SSP1 NRS1473-> DS1677 |
| BAL836 (stocked as NRS1113) |  | JH642 *amyE::Phy-spac-gfp mut2* (spc) | (Stanley et al., 2003) |
| NRS6958 | | NCIB 3610 Δ*srfAA* (kan) | (Morris et al., 2022) |
| NRS6963 | Δ*srfAA* | NCIB 3610 Δ*srfAA* (kan) *amyE::Phy-spac-gfp mut2* (cml) | SSP1 BAL834 -> NRS6958 |
| NRS1433 |  | 168 pgsB::spc | (Stanley & Lazazzera, 2005) |
| pNW654 | | Δ*motB* in pMAD | (Cairns et al., 2013) |
| NRS3494 | | 3610 Δ*motB* | pNW654 → NCIB3610 |
| NRS3434 | ΔmotB | 3610 ΔmotB +<br>*pgsB::spc* | SPP1 NRS1433 →<br>NRS3494 |

## Notes

### Competing Interest Statement

The authors have declared no competing interest.

### Summary of Updates

New title Inclusion of supplementary material

